# Extracellular vesicles promote proliferation in an animal model of regeneration

**DOI:** 10.1101/2024.03.22.586206

**Authors:** Priscilla N. Avalos, Lily L. Wong, David J. Forsthoefel

## Abstract

Extracellular vesicles (EVs) are secreted nanoparticles composed of a lipid bilayer that carry lipid, protein, and nucleic acid cargo between cells as a mode of intercellular communication. Although EVs can promote tissue repair in mammals, their roles in animals with greater regenerative capacity are not well understood. Planarian flatworms are capable of whole body regeneration due to pluripotent somatic stem cells called neoblasts that proliferate in response to injury. Here, using transmission electron microscopy, nanoparticle tracking analysis, and protein content examination, we showed that EVs enriched from the tissues of the planarian *Schmidtea mediterranea* had similar morphology and size as other eukaryotic EVs, and that these EVs carried orthologs of the conserved EV biogenesis regulators ALIX and TSG101. PKH67-labeled EVs were taken up more quickly by S/G2 neoblasts than G1 neoblasts/early progeny and differentiated cells. When injected into living planarians, EVs from regenerating tissue fragments enhanced upregulation of neoblast-associated transcripts. In addition, EV injection increased the number of *F-ara*-EdU-labelled cells by 49% as compared to buffer injection only. Our findings demonstrate that regenerating planarians produce EVs that promote stem cell proliferation, and suggest the planarian as an amenable *in vivo* model for the study of EV function during regeneration.

## Introduction

Regeneration requires the coordination of many processes including immune responses, apoptosis, cell survival, proliferation, differentiation, migration, gene expression, and extracellular matrix remodeling^1^. These processes and cell behaviors are regulated non-cell-autonomously by secreted cues like hormones^2^, cytokines^3^, and growth factors^4^. Extracellular vesicles (EVs) have recently been recognized as another mechanism for intercellular communication during regeneration^5,6^. The term "EV" broadly refers to nanoparticles with a phospholipid bilayer that are naturally secreted by cells, originating either from the endosomal trafficking system (exosomes) or by plasma membrane budding (ectosomes). EVs transport many molecular cargoes (including lipids, RNAs, and proteins) which can serve as signals to influence target cell states or behaviors. Most cells are thought to be capable of being both a donor, capable of EV production, and a recipient, capable of EV uptake^7,8^.

EVs can improve mammalian regeneration, as numerous studies have demonstrated that vesicles harvested from cultured stem cells restore pre-injury function to various tissues in damage models by enhancing processes (e.g., proliferation) integral to successful regeneration^6,9,10^. However, mammalian models used so far have a limited natural ability to regenerate, and *in vivo* studies of how tissue injury modulates EV biogenesis and function are rare. Although animals with greater regenerative abilities produce EVs^10–13^, little is known about these EVs’ roles during regeneration. Investigations in emerging animal models of regeneration could therefore complement efforts to develop EVs as tools for regenerative therapy.

The planarian flatworm *Schmidtea mediterranea* regenerates its whole body in two weeks after surgical amputation^14^. After injury, planarians develop a blastema, an unpigmented mass of differentiating cells that replace missing tissues and organs and integrate with remaining wound-proximal tissue^15^. These differentiating cells are the progeny of adult pluripotent somatic stem cells called neoblasts, which make up 20-25% of all planarian cells^16^. Neoblasts drive regeneration since they are the only somatic cell type capable of proliferation, and therefore the only source of new cells and tissue^17^. Although numerous extrinsic cues (e.g., axial polarity cues and extracellular matrix molecules) regulate specification and differentiation, only a few secreted factors are known to modulate neoblast proliferation^18^. Although EVs represent another potential class of extracellular regulators that could influence neoblasts, little is known about planarian EVs and/or their roles during regeneration. Here, we demonstrate that planarians produce EVs, and identify a likely role for EVs in promoting neoblast proliferation.

## Methods

### Animal care

Asexual *Schmidtea mediterranea* (clonal line CIW4)^19,20^ animals were maintained at 22°C in 1x Montjuic water with 50 μg/ml Gentamicin Sulfate (GeminiBio, 400-100P) in Rubbermaid containers^21^. Animals were fed every other week with calf liver paste (Sprouts Farmers Market, Oklahoma City). Water was changed one and seven days after feeding. All experiments were initiated with animals starved 6-8 days.

### Planarian dissociation

For EV isolation (below), intact animals (5-7 mm) were immobilized on filter paper cooled on a Corning CollBox XT plate, then amputated into head, trunk, and tail with a razor blade. Fragments were allowed to regenerate for 12 hr in an incubator at 22°C. Intact or regenerating animals were transferred to a microcentrifuge tube, rinsed in 1 ml of 0.22 µm-filtered CMF-B (Calcium-Magnesium Free medium plus BSA: 15 mM HEPES pH 7.4; 460 mg/l NaH_2_PO_4_ x H2O; 800 mg/l NaCl; 1200 mg/l KCl; 800 mg/l NaHCO_3_; 240 mg/l D-glucose; 1% BSA (Sigma Adrich; A3912-100G); and 50 µg/ml gentamicin) then dissociated in 0.2 ml of CMF-B plus 1 mg/ml collagenase (Sigma Aldrich; C5138-1G) per animal (e.g., 10 ml per 50 animals). Animals were gently smashed with a Kontes micropestle to start trituration, then the mixture was gently pipetted up and down with a P1000 pipet (30 strokes over 30-60 min), with gentle rocking at RT (50 rpm on a rocking shaker, Reliable Scientific, Inc. Model 55) between triturations. EV enrichment was then conducted as described below.

For cell isolation for uptake experiments or after F-*ara-*EdU labeling (below), intact or regenerating animals were placed in a 100 mm petri dish and rinsed in CMF-B, then worms were incised several times along the anterior-posterior axis with a razor blade. Animals were transferred to a microcentrifuge tube with 1 ml CMF-B plus collagenase per 2-6 animals. Tissue was triturated by pipetting up and down as above over 30-60 min, cell suspension was centrifuged at 200 x *g* for 5 min at 4°C, supernatant was removed and pellet was resuspended in CMF-B (uptake) or 1X PBS (EdU labeling) then passed through a 70 µm filter (Miltenyi Biotec MACS SmartStrainers, 130-098-462) to remove aggregates.

### EV enrichment by precipitation and size exclusion chromatography

After dissociation of 100 animals (5-7 mm in length), cell suspension in CMF-B was centrifuged at 200 x *g* for 5 min at 4 °C, and supernatant was transferred to a new tube. Cell pellet was resuspended in CMF-B, passed through a 70 µm filter, and cell density was counted in a Bio-Rad Counting Slide (Bio-Rad; 145-0011) and TC20 Automated Cell Counter (Bio-Rad). Supernatant was centrifuged at 3,000 x *g* for 30 min (unless otherwise noted, all centrifugation steps were performed at 4°C using a swing-out rotor), supernatant was again transferred to a new tube, then spun at 10,000 x *g* for 30 min. After transferring the 10,000 x *g* supernatant to a fresh tube, an equal volume of 0.22 µm-filtered 20% PEG 10,000 (Alfa Aesar B21955, dissolved in 1x PBS) was added (10% final concentration) and thoroughly mixed by inversion. The mixture was incubated for 1 hr at 4°C and centrifuged at 3,000 x *g* for 30 min to pellet EVs. Supernatant was discarded and the pellet was resuspended in 100 µl (for NTA) or 90 µl (for microinjections) of 0.22 µm-filtered isolation buffer (IB - SmartSEC Mini EV Isolation system, SBI SSEC100A-1). EVs were column-isolated exactly as per manufacturer’s instructions. The column was centrifuged at 500 x *g* for 30 sec (fixed angle rotor Eppendorf centrifuge 5424R) to collect fractions 1 (F1) and 2 (F2). For EV microinjections, 10 µl of 5 mg/ml 546 fluorescent 10,000 MW dextrans (Invitrogen, D22911) were added to F1 after SEC for a final volume of 100 µl.

### NTA

EV concentration and size were quantified using the Nanosight model NS300 (Malvern Instruments, UK – software NTA version 3.4) according to the manufacturer’s instructions. Briefly, filter 1, green laser 532 nm and SCMOS options were selected. For individual experiments the camera level and focus were adjusted and kept the same across samples. Syringe load was set at 60 µl and 3 videos of 180 sec were recorded for every tube. Sample dilutions were made right before recording to adjust concentrations. 200 nm FluoSpheres (Invitrogen, F8811) were diluted 2:73 in a final volume of 1 ml and analyzed first to verify instrument function. Lines were washed between samples with 1X PBS. Fresh SEC EVs (from 100 *egfp(RNAi)* animals resuspended in 100 µl of isolation buffer) were diluted 1:1000 and 1:250 with 1X PBS for F1 and F2, respectively.

### A280 protein quantification

10 intact animals (8-10 mm) were placed in a 100 mm petri dish, rinsed in CMF-B, diced with a razor blade, and transferred to a 15 ml tube with 10 ml of CMF-B plus collagenase. In a second tube 10 ml of CMF-B with collagenase only were aliquotted as a control. Tissue was triturated by pipetting up and down 2-3 times over 30 min, cell suspension was passed through a 70 µm filter and centrifuged at 200 x g for 5 min at 4°C (unless otherwise noted, all centrifugation steps were carried out at 4°C). The cell pellet was discarded, the supernatant (or CMF-B plus collagenase only) was transferred to new 15 ml tubes and serially centrifuged at 3,000 x *g* and 10,000 x *g* as above. The supernatant was transferred to 50 ml tubes and equal volumes (10 ml) of 20% PEG 6000 were added to reach a final concentration of 10% PEG 6000. The solution was thoroughly mixed by inverting, incubated at 4°C for 1 hr and centrifuged at 3000 x *g* for 30 min. The supernatant was removed and the pellet was resuspended in 200 µl of IB (SmartSEC Mini Kit), and split into two 100 µl aliquots. 200 µl of IB were added to the pellet-less control tube and split into two 100 µl aliquots. One tube with EVs and one control tube were further enriched with SEC as described above and only F1 was collected for measurements. Protein concentration readings were measured with a DeNovix spectrophotometer using the Protein A280 application for BSA from the nanodrop port. CMF was used to blank the instrument for readings.

### EV negative staining for TEM imaging

A drop (1-2 μl) of undiluted EVs (PEG/SEC-enriched EVs from 100 uninjured *egfp(RNAi)* animals) was placed on a formvar (EMS 15810)-coated hexagonal grid (EMS G300-Cu) for one minute, then the water in the sample was absorbed with the jagged edge of a filter paper. A drop of 2% Uranyl Acetate (EMS 22400) was layered on the sample for one minute and then absorbed. Sample was allowed to dry for one minute before grid storage at RT in a sample box. Samples were imaged on a Hitachi H-7600 transmission electron microscope equipped with an AMT NanoSprint12 at 60-80 kV. A 800 ms x 4 drift frames exposure and 5000x-25,000x magnification were used. Images were visualized and exported with AMT Capture Engine version 7.0.1.50.

### Anti-ALIX and anti-TSG101 antibody generation

Antibodies were produced by GenScript USA (Piscataway, NJ). For both ALIX and TSG101 the sequences corresponding to the full protein (dd_Smed_v6_1686_0_1 ALIX: nucleotides 62-2620, amino acids 1–852 and dd_Smed_v6_1373_1_1 TSG101: nucleotides 67-1236, amino acids 1–389) were used as the target antigen. GenScript synthesized expression constructs to generate fusion proteins, which were purified and injected into New Zealand rabbits. Antibodies were affinity purified, titered, and tested for affinity with ELISA and Western blot by Genscript.

### Protein isolation and quantification for antibody validation

For protein isolation from cells, cell pellet from 100 animals (dissociated as described above) was re-suspended in 50 ml of 1X CMF and cell density was counted. Cell suspension was split into two 25 ml equal halves, centrifuged at 200 x *g* and supernatant was discarded. For total protein quantification, one half of the cells was resuspended in 100 µl of 1X RIPA without DTT (150 mM NaCl, 1% NP-40, 0.5% sodium deoxycholate, 0.5% SDS, 50 mM Tris pH 8, and 1 µl/ml of Halt Protease inhibitor cocktail (Thermo Scientific, 78430) dissolved in water) for every 3 x 10^6^ cells. For Wes-based total protein detection and Western blot, the second cell pellet was resuspended in 100 µl of 1X RIPA with 0.1% SDS and 40 mM DTT. Cell suspensions were lysed by passing through a 25G syringe needle five times.

For protein isolation from EVs, EVs from 200 animals were enriched by PEG/SEC using two columns as described above. Each SEC fraction (100 µl x 2 columns = 200 µl for each fraction) was pooled, then split into two 100 µl aliquots. One aliquot was mixed with 100 µl of 1X RIPA without DTT and the other half was resuspended in 100 µl of 1X RIPA with 0.1% SDS and 40 mM DTT.

Both cell and EV lysates were rocked at 50 rpm for 20 min at 4°C, then centrifuged at 20,000 x *g* for 10 min at 4°C (fixed angle rotor). EV lysates were concentrated in Amicon Ultra 3 kDa columns (UFC500396) from 200 µl to 48 µl following the manufacturer’s instructions. All lysates were stored at −80°C.

Total protein was quantified using 1 ml cuvettes with a BCA Assay Kit (Thermo Scientific Pierce, 23225) according to the manufacturer’s instructions. Protein concentrations were measured using a DeNovix spectrophotometer colorimetric app for 10 mm cuvette.

### Capillary Western Blot (Wes)

Wes reagents were prepared according to the manufacturer’s instructions. Cell and Fraction 1 EV lysates were diluted to 0.5 mg/ml, and Fraction 2 EV lysates were diluted to 0.3 mg/ml. Antibodies were used at the following concentrations: custom rabbit anti-ALIX, 1.48 µg/ml; custom rabbit anti-TSG101, 22 µg/ml; custom rabbit anti-PIWI-1, 7.71 µg/ml^22^; rabbit anti-Histone H3, 20 µg/ml (Abcam, ab1791)^23^; and rabbit anti-αTubulin, 2.7 µg/ml (Abcam, ab52866)^24,25^. anti-αTubulin recognized a ∼54 kD band, close to the 50 kD predicted mass of two planarian αTubulin homologs encoded by dd_Smed_v6_38_0_1 and dd_Smed_v6_10847_0_1. Other antibodies were validated as described below. All reagents were loaded on a 12-230 kDa plate. The Separation module (SM-W008) and plate were loaded in the Simple Western (Biotechne/ProteinSimple) Wes instrument and the total protein program was selected (separation load time 200 sec, stacking load time 15 sec, sample load time 9 sec, separation time 25 min, separation voltage 375 V, biotin labeling or blocking time 30 min, primary antibody time 30 min, total protein HRP or secondary antibody time 30 min) in the Compass software (v4.0.0). High Dynamic Range Exposures were used for band detection, and contrast was manually adjusted for each antibody, but kept constant across all samples. Images of full lane labeling is provided in Source Data.

### Cloning and dsRNA feeding

*alix* and *tsg101* were amplifed from cDNA using the Bio-Rad iScript kit and cloned into pJC53.2 (RRID:Addgene_26536) as previously described^26^. For *alix,* (dd_Smed_v6_1373_1_1), nucleotides 1499-2832 were cloned (forward primer ACTCCGTCCGGGAAATTGAC; reverse primer CAATAACTGGATGCTGCGGC). For *tsg101* (dd_Smed_v6_1686_0_1), nucleotides 183-1402 were cloned (forward primer TGATGGTGTTCAGCGGACAT; reverse primer ACACACCGGCTCAATAACGT). dsRNA was synthesized as described^27^. For knockdown, dsRNA (1 μg/μl) and 1:10 blue food coloring (Durkee, B00AIYDNWA) were mixed with 2:1 liver homogenate diluted in water as described^22,26^. For *alix* RNAi, animals were fed 5 μg *alix* dsRNA per 50 μl liver homogenate, twice a week for a total of five feedings. For *tsg101* RNAi, animals were fed 1.25 μg *tsg101* dsRNA in 50 μl one time (*tsg101* RNAi was lethal 7-10 days after only one feeding). Equivalent amounts of *egfp* dsRNA^28^ were fed as a control knockdown in each group.

### Planarian-specific antibody validation

For validation of anti-ALIX specificity, 50 uninjured *egfp(RNAi)* and 50 *alix(RNAi)* animals were dissociated, then the cells for each group were divided into two equal aliquots and centrifuged. One aliquot was lysed in 200 µl of 1X RIPA without DTT and the second aliquot was lysed in 200 µl of 1X RIPA with 40 mM DTT. For anti-TSG101, 7 uninjured *egfp(RNAi)* or *tsg101(RNAi)* animals were lysed directly in 250 µl of 1X RIPA wihtout DTT and homogenized using a motorized Kontes pestle grinder. A second group of 7 uninjured *egfp(RNAi)* and *tsg101(RNAi)* animals were lysed directly in 250 µl of 1X RIPA with 40 mM DTT and homogenized. For anti-PIWI-1, 10 uninjured or 5 X-irradiated uninjured animals (7 days after irradiation) were dissociated, then the cells for each group were divided into two equal aliquots, centrifuged, and lysed in either 250 µl of 1X RIPA without DTT or 250 µl of 1X RIPA with 40 mM DTT.

Wes was performed as above, except anti-Alix was used at 3.7 µg/ml. Lysates were loaded at 0.3 mg/ml for ALIX detection, and 0.5 mg/ml for TSG101 and PIWI-1 detection. ALIX’s mass was predicted to be 95 kD and TSG101 43 kDa. Anti-ALIX labeled a prominent band by WES analysis at 98 kDa, and TSG101 at 52 kDa. Both were reduced in *alix* and *tsg101* knockdown protein lysates (Supp. Fig. 1). Similarly, anti-PIWI-1 labeling was dramatically reduced after irradiation, which ablates PIWI-1-positive neoblasts^29^.

### Planarian Irradiation

A Biological Research X-Ray irradiator (Rad Source Technologies, RS 2000) equipped with an x-ray tube was warmed up for 20 min. Animals were transferred to a 60 mm Petri dish half-filled with Monjuïc water then placed inside the irradiator. The dish was placed on top and center of the shelf, which was placed at level 5 (closest to the top of the chamber). Animals received a 60 Gy of irradiation. Animals were kept at 22°C for 1-4 days before flow cytometry analysis as controls for setting X1 and X2 gates or 7 days for Protein isolation and WES detection for anti-PIWI-1 antibody validation.

### EV enrichment and PKH67 labeling of EVs for uptake assay

360 intact animals (6-7 mm) were dissociated for EV enrichment as follows. Worms were placed in a 100 mm petri dish, rinsed in CMF-B and diced with a razor blade. Animals were transferred to three 50 ml tubes with 25 ml of CMF-B plus collagenase each. Tissue was triturated by pipetting up and down over 30-60 min, cell suspension was passed through a 70 µm filter and centrifuged at 200 x *g* for 5 min at 4°C (unless otherwise noted, all centrifugation steps were carried out at 4°C). The supernatant was transferred to clean 50 ml tubes, centrifuged at 3,000 x *g* for 30 min and then at 10,000 x *g* for 30 min. The supernatant was divided into three 25 ml aliquots and 25 ml of 16% PEG 6,000 (Alfa Aesar; A17541, dissolved in 1X PBS)^30^ was added to each aliquot to reach a final concentration of 8% PEG 6,000. The solution was mixed, incubated overnight at 4°C and centrifuged at 3,000 x *g* for 1 hr. The supernatant was discarded and EV pellets were resuspended and pooled in a total of 9 ml of 1X PBS. 9 ml 1X PBS was also added to control solution tube, which had no visible post-precipitation pellet. 4.8 ml of EVs or control solution were mixed with 2.4 ml of 1:50 PKH67 plus Diluent C staining solution (Sigma PKH67GL-1KT)^31^. The remaining 4.2 ml of EVs were left unstained. EV and control samples were incubated in the dark for 5 min without rocking at RT. Each sample was transferred to a 50 ml tube, brought to 25 ml with 1X PBS, mixed with 25 ml of 10% PEG 6000, and incubated overnight at 4°C. Tubes were centrifuged at 3,000 x *g* for 1 hr and supernatant was discarded. The pellet of PKH67 labeled EVs was resuspended in 4 ml of CMF-B and unstained EVs in 3.5 ml (24 animals’ EVs per 0.5 ml for addition to cells, below). 4 ml of CMF-B were added to the pellet-less control solution (no EVs).

### Isolation of cells for EV uptake assay

65 intact animals (6-7 mm) were rinsed in CMF-B and diced with a razor blade, then dissociated in 60 ml of CMF-B plus collagenase. Tissue was triturated by pipetting up and down 3-4 times over 30-60 min, cell suspension was passed through a 70 µm filter and centrifuged at 200 x *g* for 5 min at 4°C (unless otherwise noted, all centrifugation steps were carried out at 4°C). The supernatant was discarded, the cell pellet was resuspended in 50 ml of CMF-B, and strained through a 30 µm nylon mesh (Miltenyibiotech MACS SmartStrainer; 130-098-458) into a new 50 ml tube. Cell densities were determined, then 6 x 10^6^ cells were aliquoted to 1.7 ml tubes, centrifuged at 200 x g, supernatant was discarded prior to cell resuspension in EVs.

### EV uptake assay by flow cytometry

Cells were resuspended in 24 animals’ worth of EVs: either 0.5 ml PKH67-labeled EVs or 0.5 ml unlabeled EVs in CMF-B, or 0.5 ml control (no EV) solution. After resuspension, cells were incubated for 0, 1, 2, or 6 hr protected from the light and while rotating at 24 rpm on a GyroMini nutating mixer (Labnet, S0500) at RT. 45 min before the incubation with EVs was done, 50 µg/ml of Hoechst 33342 were added to label nuclei. Cells were centrifuged at 200 x *g* for 5 min and resuspended in 0.5 ml of CMF-B with Hoechst and propidium iodide (PI, 1 µg/ml). 0 hr samples were processed with 2 hr samples, but cells were centrifuged immediately after resuspension to remove EVs, then incubated 1X CMF-B only until Hoechst addition. Control samples for flow cytometry (unstained, unstained cells 2 hr after PKH67-labeled EV uptake, Hoechst, PI only, double stained, and x-irradiated) were prepared in parallel with the experimental samples.

### EV uptake assay by microscopy

EVs were prepared similarly as in the uptake section above but with the following exceptions. EVs were isolated from 25 dissociated animals, and cells were isolated from 75 dissociated animals. EV pellets were resuspended in diluted Schneider’s *Drosophila* medium ("dSchneider’s," 2.8x Schneider’s *Drosophila* medium pH 7.4 (Gibco, 21720024) diluted in water plus 1x Penicillin-Streptomycin (Gibco, 15070063))^32^. dSchenider’s was also added to the pellet-less CMF-B, PKH67-labeled sample. No unstained EVs were prepared. Cells were centrifuged, resuspended, and strained one additional time through a 10 µm mesh (pluriSelect pluriStainer; 43-50010-01). 1×10^6^ cells were aliquotted per sample. Cells were incubated with EVs for 0 or 2 hr only and every 0.5 ml of dSchneider’s contained 2 animals’ worth of EVs. Nuclei were also stained with 10 µg/ml of Hoechst 33342 and no PI staining was performed. After incubation, cells were transferred to a 4-well 60 mm dish (Greiner BIO-ONE, 627870), allowed to settle for 10 min, then imaged with Zeiss AxioObserver.Z1.

### Flow cytometry for live cells

After dissociation (above) of isolated tail tissue (posterior to the pharynx and minus 1 mm of the tail tip) from 4-5 mm worms, cells were analyzed on a FACSCelesta (BD Biosciences) equipped with 405 nm, 488 nm, and 640 nm lasers and a 180 µm flow cell opening running at slow speed. Unstained, Hoechst-only, and PI-only controls were used to set PMT voltages, and X gates^33^ were set as previously described^22^: X1 (∼10% of Hoechst+, PI-events), X2 (∼20% of live cells), and Xins (∼60%). In the BD FACSDiva software version 9.0, the FSC, SSC, BV421, BV650, and PI lasers were selected.

### Live cell imaging for EV uptake

Live cells for uptake experiments were imaged using a Zeiss AxioObserver.Z1 running Zen Blue version 2.3.64.0. A 63X oil immersion objective (Plan-Apochromat 63X/1.4 NA), Hoechst (DAPI), PKH67 (GFP), and transmitted light (TL) channels were used. The same exposures were maintained across samples. 2×2 tiles were stitched post-acquisition.

### Microinjections

A Warner Instruments microinjector (Pico-Liter Injector Model PLI-90A), PicoNozzle needle mount (World Precision Instruments, Inc; 5430-ALL), and Zeiss Stemi 508 were used for microinjections. Pressure to Pico-Liter Warner Injector was set at 40 psi, injection time at 0.20 sec, pressure meter at 1 psi, and pressure balance at - 0.2 psi. Borosilicate glass capillaries (Sutter Instrument; B100-50-10) were pulled into needles on a needle puller (Sutter instruments P1000), using 563 for heat, 60 for pull, 80 for velocity, 90 for delay, 200 for pressure, 553 for ramp with delay mode and sufe heat settings on. Each needle was loaded with ∼10 µl of an EV F1/dextran mixture (above, EVs from 1 animal per µl, approximately 4×10^11^ EVs total in each 100 µl EV preparation esimated from NTA experiments with identical animal numbers) or buffer/dextran control solution (no EVs). One animal at a time was poked with a fine blade (FST; 10055-12) between the tail branches on the dorsal side, then injected with ∼1 µL (approximately 4×10^9^ EVs) for RNAseq (larger animals, 6-7 mm) and ∼0.25 µL (approximately 1 x10^9^ EVs) for anti-pH3-PS10 and EdU labeling (smaller animals, 4-5 mm). Animals were allowed to recover for at least one hour then worms without fluorescence were removed.

### RNA sequencing

Twenty-four hours after microinjection, tail fragments from 6-7.5 mm worms were amputated and ∼10% of the posterior-most tail tip was removed, then tissue was immediately homogenized in Trizol (Thermo Fisher Scientific 15596026) with a motorized Kontes pestle grinder. Three fragments per biological replicate were pooled, for a total of four biological replicates per condition (uninjected, buffer-injected, or EV-injected). RNA was extracted with two chloroform extractions and high-salt precipitation buffer as per the manufacturer’s protocol. High salt solutions were transferred to Zymo RNA columns (Zymo RNA Clean & Concentrator 5 kit, R1013) for DNAse treatment and purification as per the manufacturer’s instructions.

mRNA was enriched using the NEB NEBNext Poly(A) mRNA Magnetic Isolation module, and libraries were generated using the IDT xGen RNA Library Prep Kit according to the manufacturer’s protocol. Final libraries were assayed on the Agilent TapeStation for appropriate size and quantity. Libraries were pooled in equimolar amounts using the IDT Normalase module, then final pools were absolutely quantified using qPCR on a Roche LightCycler 480 with NEB Illumina Library Quantification Reagents. Paired-end (2□×□150Lbp) sequence was generated on an Illumina NovaSeq X Plus instrument.

BBDuk (v36.99) (https://sourceforge.net/projects/bbmap/) was used to trim paired-end reads with settings k=13 ktrim=r mink=11 qtrim=rl trimq=10 minlength=35 tbo tpe. Read quality was assessed before and after trimming with FastQC (v0.11.5)^34^. Reads were mapped to unique dd_Smed_v6 transcripts^26,35^ in Bowtie2 (v2.3.1) with “-a” multi-mapping and “–local” soft-clipping allowed. The “featureCounts” utility in the Subread package (v1.6.3)^36^ was used with a custom “.SAF” file and options “-p -M -O -F SAF” to include multi-mapping and multi-overlapping reads for read count summarization.

Read counts were analyzed in edgeR v3.26.3^37^. First, all transcripts with counts per million (CPM)□<□1 in less than four samples (e.g., lowly expressed transcripts) were excluded from further analysis; 17,755 transcripts were retained after this cutoff. Library sizes were recalculated, then samples were normalized using the trimmed mean of M-values (TMM) method, and generalized linear model (GLM) common, trended, and tagwise dispersions were calculated. The negative binomial GLM was fitted with glmQLFit, followed by identification of differentially expressed transcripts were identified using the GLM quasi-likelihood ratio test (glmQLFTest). Expression changes were considered to be significant if the false discovery rate-adjusted *p* value (“FDR”) was <0.05.

Cross-referencing with transcripts enriched in X1, X2, Xins, PIWI-1-high, PIWI-1-low, and PIWI-1-negative subpopulations was performed as previously described^22^ (https://zenodo.org/records/6596520). For cross-referencing of EV-responsive transcripts and injury-responsive transcripts in blastemas, differential expression data from published bulk RNA-Seq data were used^38^. Specifically, logFCs and adjusted *p* values from 6 hr vs 0 hr, 24 hr vs 0 hr, and 48 hr vs 0 hr comparisons in control (*unc-22*) RNAi samples from the *myoD* RNAi experiment were compared to RNA-Seq data in this study.

### Anti-phospho-histone H3-PS10 immunolabeling

12 and 24 hpi, mucus from injected animals was removed by cooling worms on ice for 1 min then incubating on ice-cold 2% HCl (Sigma H1758) for 3 min. Samples were fixed in methacarn (60% MeOH (Fisher Scientific A4124), 30% Chloroform (Fisher Scientific C298), and 10% Acetic acid (Fisher Scientific A38-212)) for 20 min. Animals were washed twice in methanol then stored at −30°C. Fixed worms were bleached in 5% H_2_O_2_ in MeOH and placed 5-10 cm under direct fluorescent light for 15 hr, then rinsed 3x in MeOH. Fixed animals were amputated in half, then gradually rehydrated in methanol and PBS-Tx (75% methanol/25% PBS-Tx, then 50%/50%, then 25%/75%) then incubated in blocking solution (0.3% Triton-X100, 0.45% fish gelatin (Sigma G7765-250ML), 0.6% IgG free BSA (Jackson ImmunoResearch 001-000-162) dissolved in 1xPBS) for 4 hr. Fragments were incubated in 1:2000 rabbit anti-PS10 (Cell Signaling #3377S, clone D2C8) in blocking solution for 15 hr at 4°C. Animals were washed in PBS-Tx for 8 times, 30 min per wash. Worms were re-blocked for 1 hr then incubated in 1:2000 Goat anti-Rabbit-HRP (Jackson ImmunoResearch 111-035-003) solution for 15 hr at 4°C. Worms were washed in PBS-Tx for 8 x 30 min each. Tyramide signal amplification (TSA) was conducted as described^39^. Samples were rinsed 4 times then washed for 15 hr at 4°C. Animals were incubated in Vectashield for at least 2 hr at 4°C before mounting.

### Imaging and quantification for pH3-PS10 immunolabeling

For imaging of anti-pH3-PS10 immunolabeled samples, z-stacks of mounted animals (tail region, dorsal side up) were collected on a Leica M205 FCA Fluorescence stereo microscope. The magnification was set at 8.04X, DAPI (ET DAPI) and pH3-PS10 (ET mCHER) channels we used. The same exposures were maintained across samples. After image acquisition parallax correction was applied in the LAS X software (version 3.4.1.17822). Animal area and number of PS10+ cells were quantified in FIJI running ImageJ 1.53t^40^. Briefly, a z-projection was created from the parallax files by selecting the max intensity for projection type, and the channels were split. "Auto Local Threshold" with method Otsu and default settings was used on the DAPI image to identify the animal area, which was outlined using the wand tool, manually cropped to the post-pharyngeal region if necessary, then measured. pH3-PS10+ cells were counted using the "Find Maxima" function within the outline generated with the DAPI channel. The area and pH3-PS10 count results were exported to excel where the number of mitosis per mm^2^ were calculated.

### F-*ara*-EdU labeling

1 hr after microinjection, mucus was removed by rinsing animals (sized between 4-5 mm) in 0.0625% N-Ac for 1 min. The worms were rinsed 3 times in Montjuïc water and transferred to a well (20 worms per well) in a 24-well plate. Planarians were soaked in 1 ml of 0.5 mg/ml F-*ara*-EdU (Sigma-Aldrich, T511293-5MG – plus 3% DMSO dissolved in Montjuïc water) and kept protected from light. Animals were allowed to soak for 11 hr at 22°C then analyzed by flow cytometry.

### F-*ara-*EdU detection and flow cytometry of fixed cells

After dissociation (above) of isolated tail tissue (posterior to the pharynx and minus 1 mm of the tail tip) from 6-7 mm worms, cells were centrifuged (unless otherwise noted, all centrifugation steps were performed at 4°C for 5 min using a swing-out rotor) at 300 x *g*, re-suspended in ice cold 1X PBS and strained through a 70 µm nylon mesh. Cells were fixed for 10 min (unless otherwise noted, incubations and washes were performed at RT while rocking at 50 rpm) in 1.92% Paraformaldehyde (PFA – Ted Pella, Inc. 18505), centrifuged (from this point onwards all centrifugations were at 600 x *g*), resuspended in 1X PBS, and stored at 4°C until all samples were fixed. Samples were centrifuged, re-suspended in 0.3% PBS-Tx (1X PBS and 0.3% Triton-X100), permeabilized for 30 min, centrifuged and re-suspended in 3% PBS-B (1x PBS and 3% BSA). The tubes were centrifuged, the supernatant discarded and the cells re-suspended in 500 µl of azide labeling solution (1X PBS, 1 mM CuSO_4_ (Sigma-Aldrich, C1297-100G), 100 µM Fluorescein Azide (Sigma-Aldrich, 910147-10MG), and 100 mM (+)-Sodium L-Ascorbate (Sigma, A7631-100G)) and incubated for 30 min. Cells were washed once in 3% PBS-B, centrifuged, re-suspended in 3 µg/ml DAPI to label nuclei for 10 min, centrifuged, resuspended in 175 µL of 1X PBS with 0.5% BSA per tube, and filtered through a 40 µm mesh into a flow cytometry tube. Cells were analyzed on a FACSCelesta running at slow speed. Unstained, DAPI-only, and EdU-only controls were used to set PMT voltages. BV421, Alexa Fluor 488, and BV711 (autofluorescence) channels were used.

### Statistics

For experiments with statistical analysis, the values for n (indicated by the number of data points in figure plots), error bars and *p* values are found in legends. Tests were conducted in Prism 9 (GraphPad Software, San Diego, CA). Brown-Forsythe and Welch ANOVA tests were performed with Dunnett’s multiple comparisons correction. Unpaired t-test was performed with Mann Whitney correction. Asterisks in figures indicate *p* values of <0.05 (*), <0.01 (**), <0.001 (***), and <0.0001 (****). For comparison of transcript numbers responsive to EV or buffer injection, Fisher’s exact test was performed by comparing the number of signature transcripts in each gate that were up- or down-regulated vs. remaining unchanged transcripts in that gate, for buffer-injected vs. uninjured and EV-injected vs. uninjured conditions. For more details about statistical testing see Source Data.

### Data Availability

Processed RNA-Seq data associated with this study are provided in Supplementary Table 2, and will made publicly available in the NCBI Gene Expression Omnibus (GEO) upon acceptance of manuscript. Other data supporting this study’s findings are available within the article and its Supplementary files, or from the authors upon reasonable request.

### Ethics Statement

Antibodies were produced by GenScript USA (Piscataway, NJ), an OLAW, AAALAC, and PHS-approved vendor. GenScript’s animal welfare protocols were approved by Oklahoma Medical Research Foundation IACUC (17–58). No other vertebrate animals were used in this work.

## Results

### Planarians produce extracellular vesicles

To determine whether planarians produce EVs, we used the most widely adopted method to dissociate uninjured planarians into single-cell suspensions: manual dicing of tissue fragments, then gentle trituration in calcium- and magnesium-free medium with bovine serum albumin and collagenase to disrupt the extracellular matrix (ECM)^33^. After serial centrifugation at 200 x *g,* 3,000 x *g,* and 10,000 *x g* to remove cells and cellular debris, we enriched EVs by precipitation with the polymer polyethylene glycol (PEG, 10,000 kDa)^30,41–43^ followed by resuspension and size exclusion chromatography (SEC) rather than ultracentrifugation, to better maintain EV activity and minimize damage^44–47^ (Fig. 1A). We used Nanoparticle Tracking Analysis (NTA) to measure size distribution and concentration of EVs. The average mean nanoparticle size was 152.55 nm for fraction 1 (F1) and 124.35 nm for fraction 2 (F2), with overall sizes ranging between 30 and ∼300 nm, consistent with published size ranges for EVs from many sources (Fig. 1B)^48–50^. We did not analyze fraction 3, as total protein content was negligible (not shown). Normalized particle count was ∼4,700 and 965 particles per cell for F1 and F2, respectively (Fig. 1C). PEG and PEG/SEC eluates from media alone had very low protein content (Supp. Table 1), demonstrating that media components (BSA and collagenase) were not co-isolated by PEG precipitation or SEC. We then used transmission electron microscopy (TEM) to analyze the SEC eluate, and observed nanoparticles with a cup-like shape, similar to EVs from other organisms^51,52^, including other invertebrates^13,53,54^ (Fig. 1D).

**Figure 1.**
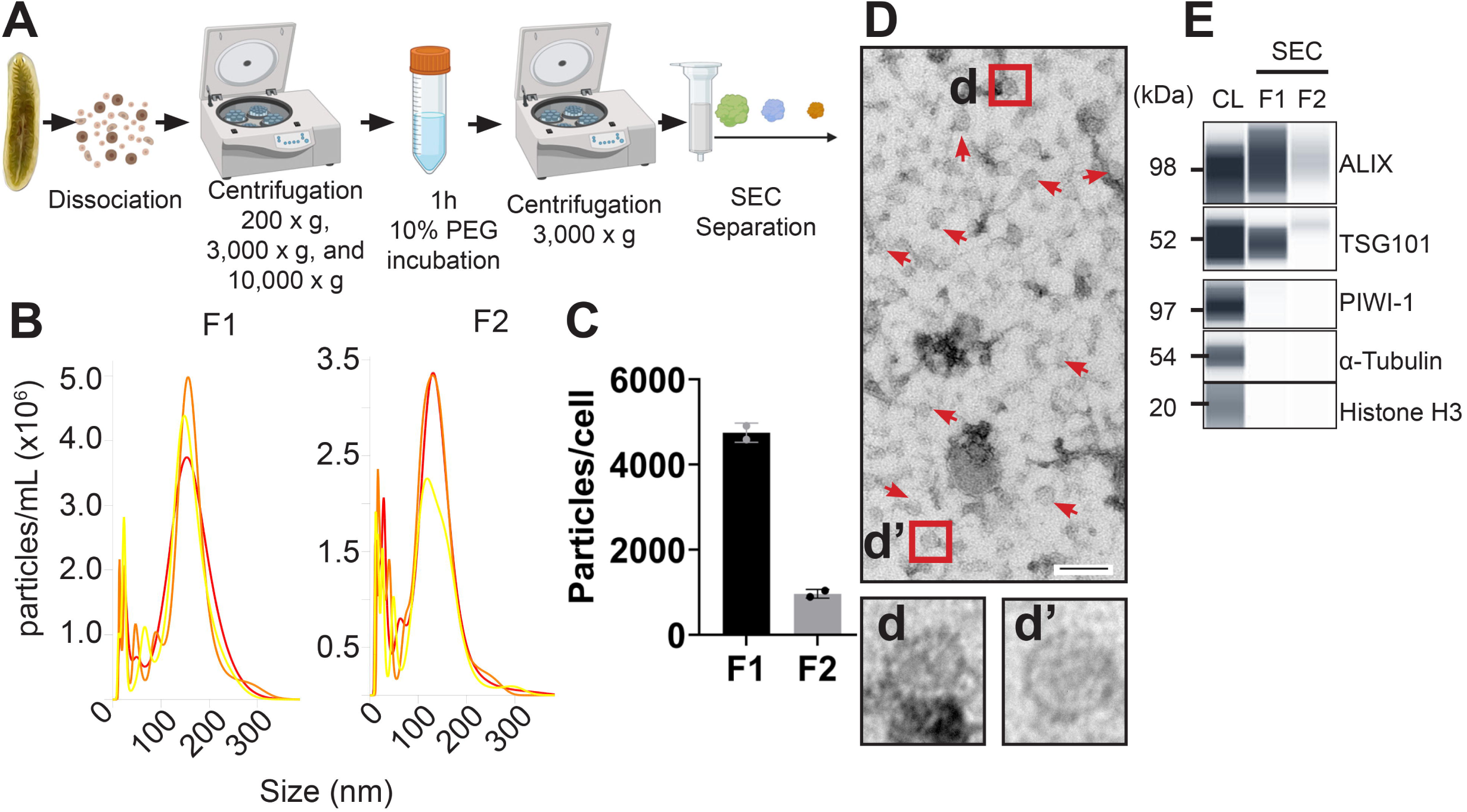
Planarians produce extracellular vesicles. **(A)** Schematic of EV enrichment by sequential centrifugation followed by 10% PEG 10,000 precipitation and SEC. **(B)** Representative curves of size distribution analysis performed with NTA. **(C)** Particle number normalized to cell count showing two technical replicates. **(D)** Transmission electron micrograph of EVs (red arrows). **(d-d’)** Insets show enlargement of the boxed regions. **(E)** Simple Western (capillary-based Western blot) of EV markers ALIX, TSG101 and non-EV markers PIWI-1 (cytoplasmic), α-Tubulin (cytoplasmic), and Histone H3 (nuclear). Cell or EV lysates were loaded at equal protein concentrations for all antibodies. Scale bars: 100 nm (**D**). ALIX, ALG-2-interacting protein X; TSG101, Tumor suppressor gene 101; PIWI-1, P-element-Induced WImpy testis-1; SEC, size exclusion chromatography; CL, cell lysate; F1 and 2, Fraction 1 and 2. Panel A created with Biorender.com.

To address whether the planarian nanoparticles isolated were bona fide EVs, we isolated protein and performed Simple Western analysis (a size-based capillary Western blot assay), to assess protein content of the first two SEC fractions. We also generated custom polyclonal antibodies against planarian orthologs of two of the most commonly used EV markers, the ESCRT complex accessory protein ALG-2-interacting protein X (ALIX) and ESCRT-1 complex component Tumor Susceptibility Gene 101 (TSG101)^55–57^, as well as PIWI-1, a cytosolic RNA-binding protein expressed by neoblasts and their early progeny^22,29^ (Supp. Fig. 1A-C). Planarian nanoparticles carried both ALIX and TSG101, but not PIWI-1 (Fig. 1E). In addition, using antibodies that cross-react with planarian proteins, we also found that enriched EVs did not carry the cytoskeleton component α-Tubulin^24,25^ or the nuclear protein histone H3^58^ (Fig. 1E). Together, these data show that precipitation combined with SEC selectively enriches EVs and not cellular debris generated by tissue dissociation, and indicate that planarians produce EVs.

### Planarian cells take up EVs

Since EVs exert their influence on gene expression and cell behaviors through direct interaction with target cells^59^, we next tested whether neoblasts could take up EVs. We enriched planarian EVs using PEG precipitation, labeled EVs with the lipophilic dye PKH67, which labels the membrane of EVs^60^, then incubated these labeled EVs with dissociated cells (Fig. 2). As negative controls, we also incubated cells with either unlabeled EVs, or PEG precipitate prepared from an identical volume of media containing the same concentration of PKH67 used to label EVs (Supp. Fig. 2). Using live cell flow cytometry (see gating strategy and controls in Supp. Fig. 2A-E), we quantified PKH67^+^ cells in three subpopulations distinguished by Hoechst 33342 blue (DNA content) and Hoechst red fluorescence^33,61^ (Supp. Fig. 2D). The X1 gate captures radiosensitive neoblasts in S and G2/M phases of the cell cycle, which are ablated 24 hours after a lethal dose of X-irradiation (60 Gy) (Fig. 2A and Supp. Fig. 2D-E)^61,62^. The X2 gate is composed of a mixture of G1 neoblasts and G0 post-mitotic early progeny, which are lost more gradually after X-irradiation^63^. Cells in the Xins (X-insensitive) gate consist of irradiation-insensitive later progeny and fully differentiated cells. At 1, 2 and 6 hours post culture an increasing percentage of X1, X2, and Xins cells all became PKH67^+^ (Fig. 2B-E). A proportion of cells in all three gates showed increased green fluorescence as soon as one hour after incubation (Fig. 2B-E). By 6 hr, ∼42% of X2 cells and ∼58% of Xins cells had become PKH67^+^ (Fig. 2B, middle and right, and Fig. 2D-E). By contrast, ∼98% of X1 neoblasts had become PKH67^+^ (Fig. 2B, left, and Fig. 2C).

**Figure 2.**
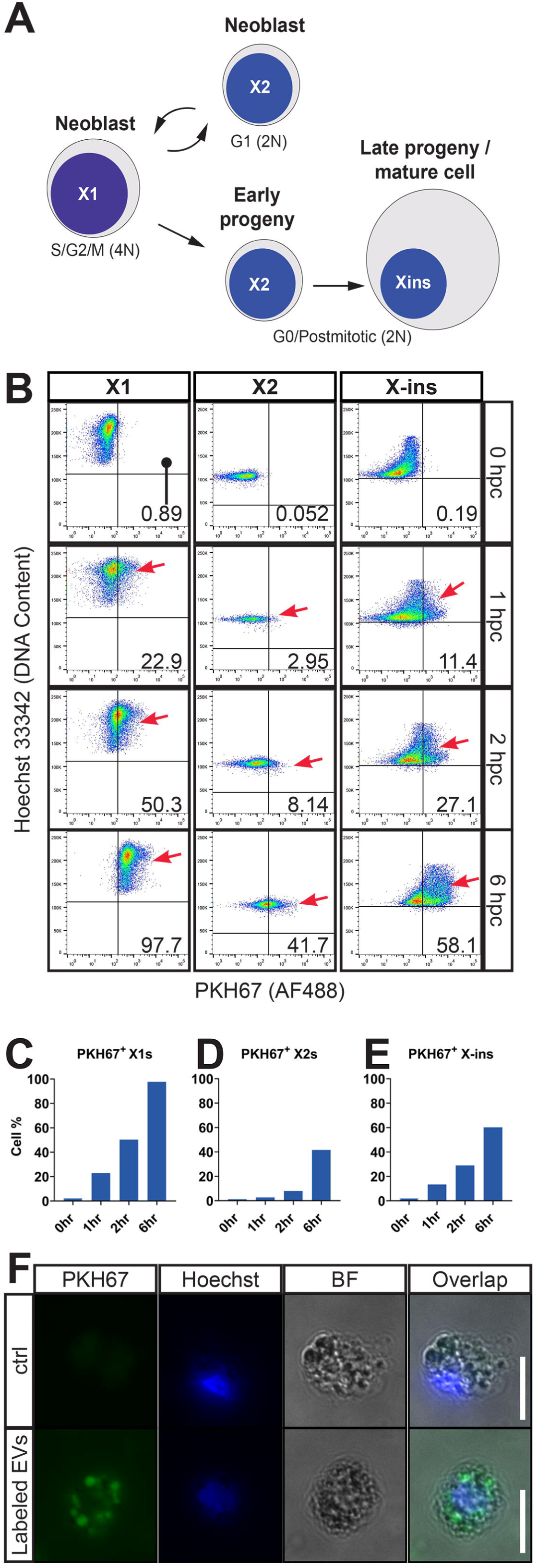
EVs are taken up by planarian cells. **(A)** Schematic representation of flow cytometric cell subpopulations relative to DNA content and cell cycle state **(B)** Representative live cell flow cytometry plots showing the fluorescence intensity of PKH67 (Alexa Fluor 488, x-axis) of X1 (left), X2 (middle), and X-insensitive (right) cells over time. Red arrows point to the PKH67^+^ cell fractions. **(C-E)** Quantification of PKH67^+^ cells at increasing time points in the X1 **(C)**, X2 **(D)**, and X-ins **(E)** gates. **(F)** Microscopy images of live planarian cells after 2 hours of incubation with buffer (top) or PKH67 labeled EVs (bottom). Scale bars: 10 µm (**E**). hpc, hours post-culture; BF, brightfield.

Cells incubated with unstained EVs (Supp. Fig. 2F and H-J) or PEG precipitate prepared from PKH67-containing CMF media (Supp. Fig. 2G and H-J) had a negligible shift in fluorescence over time. These results suggested that unstained EVs did not increase autofluorescence of cells, and that control PEG precipitate (dye mixed with buffer) did not contain significant levels of PKH67 micelles that could result in false positive signals^64^. Qualitatively, planarian cells after two hours of incubation with labeled EVs, but not control PEG precipitate, showed obvious PKH67 fluorescence in intracellular compartments (Fig. 2F). We conclude that planarian cells bind and take up EVs in a time-dependent manner, and that different cell types and states have different binding and/or uptake rates. Furthermore, the fact that S/G2 neoblasts (X1 cells) were almost all labeled suggested that neoblast gene expression and/or behavior could be influenced by EVs *in vivo*.

### EVs modulate neoblast gene expression and promote cell proliferation

EV uptake often influences gene expression by target cells^65^. Accordingly, to test the biological activity of EVs *in vivo,* we first asked whether EVs modulated expression of transcripts associated with neoblasts, progeny, and mature cells. Reasoning that EVs isolated from regenerating tissue might promote pro-regenerative gene expression, we isolated EVs from head, trunk, and tail fragments 12 hours after amputation (hpa), a time point that slightly precedes localized proliferation and differentiation during regeneration^66,67^. Next, we injected these EVs (or buffer only) into uninjured live planarians between the intestinal branches in the tail, a region where many neoblasts reside^68^ (Fig. 3A). We then extracted RNA from tissue near the injection site 24 hours post-injection (hpi), and conducted bulk RNA-Seq (Fig. 3B, Supp. Table 2). Only a small number of transcripts (10 up and 14 down) were significantly and differentially expressed between EV-injected and buffer-injected tissue (Supp. Fig. 3A). However, ∼4700 (2556 up and 2138 down) transcripts were significantly up- or downregulated in buffer-injected samples vs. uninjected samples (Supp. Fig. 3B), while ∼5500 (2883 up and 2616 down) transcripts were differentially expressed (DE) in EV-injected vs. uninjected samples (Supp. Fig. 3C). This suggested that EV injection induced subtle, but broader changes in gene expression than buffer injection only.

**Figure 3.**
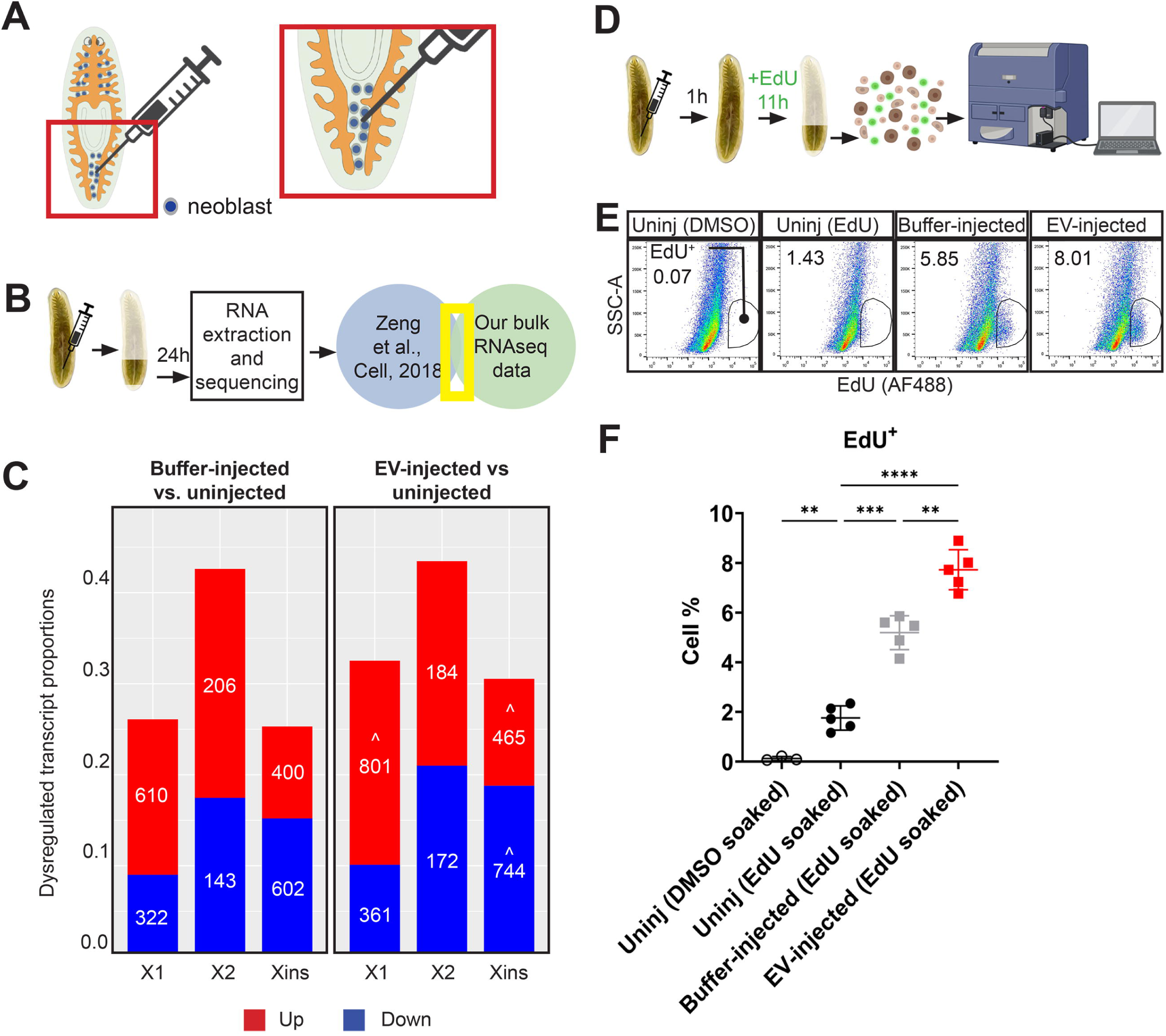
EVs increase injury-responsive gene expression changes and promote proliferation. **(A)** Schematic of EV injection into a live worm between the two posterior intestinal branches where neoblasts are more abundant. **(B)** Schematic of experimental design for bulk RNA-seq analysis **(C)**: injection of EVs from 12 hr regenerates, RNA extraction, and cross-referencing of transcript-enrichment in subpopulations of planarian cells. **(C)** Number and ratio of X1 (3,572), X2 (819), and X-ins (3,959) signature transcripts that were up- or down-regulated in buffer-injected vs. uninjured (left) or EV-injected vs. uninjured (right) animals. Statistically significant differences in transcript number for EV-injected vs. uninjected compared to buffer-injected vs uninjected are denoted with a caret (^) on the right plot (Fisher’s exact test). **(D)** Schematic of experimental design for (**E-F**): injection of EVs from 12 hr regenerates, F-*ara-*EdU soaking for 11 h, and dissociation of injected tail region. **(E)** Representative flow cytometry plots of cells dissociated from tail regions 12 hpi. **(F)** Quantification of (**E**). Brown-Forsythe and Welch ANOVA test with Dunnett’s T3 multiple comparisons test. Error bars: mean ± S.D., n = 3 (Intact DMSO), 5 (Intact EdU), 5 (buffer-injected), or 5 (EV-injected) biological replicates per condition. (**D**) From left to right **p = 0.008, ***p = 0.0002, **p =0.0036, ****p < 0.0001. Uninj, uninjected. Panels A, B, and D created in Biorender.com.

To determine whether injection affected specific cell types or states, we cross-referenced these DE transcripts with previously published lists of "signature" transcripts enriched in X1, X2, and Xins cells^22,69^ (Supp. Table 3A-C). Buffer injection caused downregulation of ∼9-17% of signature transcripts in each subpopulation, and upregulation of ∼17% of X1-enriched transcripts, ∼25% of X2-enriched transcripts, and ∼10% of Xins-enriched transcripts (Fig. 3C). EV injection showed similar trends, but more transcripts were upregulated and downregulated (Fig. 3C). For example, 610 X1 transcripts were upregulated in control-injected vs. uninjured samples, while 801 X1 signature transcripts were upregulated in EV-injected vs. uninjured samples, an ∼31% increase. We also cross-referenced our results with transcripts enriched in cells expressing the pan-neoblast marker PIWI-1, which is found in high levels in neoblasts, low levels in early progeny, and is turned off in differentiated cells^29,70^ (Supp. Table 4A-C). We observed a similar pattern of up- and downregulated genes in each subpopulation (Supp. Fig.3D). The increased number of EV-responsive genes (vs. injection/buffer-responsive) was statistically significant for upregulated X1- and PIWI-1- high-enriched transcripts, and both up- and down-regulated Xins and PIWI-1-negative transcripts (Fig. 3C and Supp. Fig. 3D). Many of the neoblast-enriched transcripts (e.g., X1 and PIWI-1-high) that were uniquely upregulated in EV-injected vs uninjured but not control-injected vs uninjured animals were also injury-responsive in bulk RNA-Seq of blastema tissue^38^ (Supp. Fig. 3E-I, and Supp. Table 5A-B). This suggested that buffer injection activated gene expression programs similar to amputation injury, and that EV injection augmented these injury-responsive expression changes. In addition, the upregulation of greater numbers of X1- and PIWI-1-high-enriched neoblast transcripts by EV injection suggested the possibility that EVs could promote or sustain neoblast proliferation.

To test this hypothesis, we immunolabeled and quantified mitotic cells after injection of EVs from 12 hpa fragments (Supp. Fig. 4A-D). In planarians, minor injuries like needle pokes stimulate increases in the number of mitotic phospho-Histone H3- S10-positive (PS10^+^) cells for up to 48 hours, indicating that minor injuries promote neoblast cell cycle acceleration and proliferation, even without tissue loss^66^. Unexpectedly, the number of PS10^+^ mitoses (per mm^2^) in EV-injected and buffer-injected animals was similar at both 12 hpi and 24 hpi (Supp. Fig. 4C-D). However, EVs could promote a transient acceleration of M-phase, reducing the number of PS10+ cells, or peaks in the number of mitotic cells might occur earlier or later than the time points we assessed.

We therefore chose to address whether EVs promoted proliferation using sustained F-*ara-*EdU labeling^58,71^ as a more cumulative method to measure proliferation and cell cycle progression (Fig. 3D-F). We injected EVs from 12 hr regenerates (or buffer only) into planarians, waited 1 hour for the wound to close, soaked animals in F- *ara-*EdU for 11 hours, and then quantified EdU^+^ cells at 12 hpi by flow cytometry (see gating strategy and controls in Supp. Fig. 5A-C). Both buffer- and EV-injected groups had higher percentages of EdU^+^ cells than the uninjured control group. However, planarians injected with EVs had ∼49% more EdU^+^ cells (Fig. 3E and F) than buffer-injected animals (7.72% vs 5.19% of DAPI^+^ cells). This result was consistent with our bulk RNA-Seq data, and strongly suggested that EV injection boosted the neoblast proliferative response vs. buffer injection alone. The majority of EdU^+^ cells had 2N-4N (S) and 4N (G2/M) DNA content (Supp. Fig. 5D and 5F-G). In EV-injected samples, 2N- 4N (S-phase) cells increased ∼50%, and 4N (G2/M) increased ∼40% compared to buffer-injected samples (Supp. Fig. 5D and 5F-G). A smaller percentage of 2N (G1/G0) cells were also EdU^+^ (Supp. Fig. 5D and E), indicating that some EdU-labeled cells underwent mitosis. Although the differences in 2N, EdU^+^ cells between EV-injected, buffer-injected, and uninjected groups were not statistically significant, the trend was higher in EV-injected animals (Supp. Fig. 5E). Together, these data suggest that EVs promote proliferation.

## Discussion

In this work, we have demonstrated that planarian flatworms produce EVs with morphology and sizes similar to other organisms. These EVs carry orthologs of EV markers but not cytosolic or nuclear exclusion markers. Planarian cells bind and likely take up EVs at different rates depending on cell cycle stage and differentiation status. Finally, planarian EVs are biologically functional *in vivo*, and can promote upregulation of injury-responsive neoblast-enriched transcripts and increase proliferation.

Regeneration requires the formation of new tissue, which usually requires an increase in cell proliferation^72^. Identification of secreted cues that regulate proliferation after tissue injury is therefore a key goal. In planarians, only a few such cues have been identified^18,73,74^. For example, knockdown of the secreted TGF-β inhibitor Follistatin results in a dramatic reduction in stem cell proliferation with a concomitant regeneration failure^75^. In addition, extracellular matrix components, hedgehog pathway signaling, and a few other regulators promote and/or inhibit proliferation^73,76–78^. Here we identify EVs as another potential mechanism by which pro-proliferative signals could be secreted and delivered to neoblasts.

EVs promote proliferation in a variety of contexts, although studies of this activity and identification of specific mechanisms during regeneration are limited. For example, the number of EVs released increases after ischemic injury in the mammalian liver and zebrafish endothelial cells^79,80^. EVs promote proliferation of mammalian cardiomyocytes, hepatocytes, and intestinal cells by transporting specific cargo (e.g., periostin, circular RNA Whsc1, or sphingosine kinase 2) to activate several pathways (e.g., Yes-associated protein Yap, STAT3/cyclinB2, sphingosine-1-phosphate, or Wnt signaling) *in vitro* and/or *in vivo*^79,81–83^. EVs also promote proliferation through Erk and Wnt signaling in zebrafish, and promote head regeneration and upregulate Wnt signaling in *Hydra* ^13,84,85^. An important next step will be to identify pro-proliferative cargo molecules and the pathways they regulate in planarians.

It will also be important to identify the cellular mechanism by which EVs promote proliferation. Previous work in planarians suggested that signals originating from even small wounds like needle pokes induce a mitotic peak between 4 and 10 hours after injury^66^. In that study, the authors proposed that the number of mitotic cells increased rapidly by shortening the G2/M phases and possibly S phase^66^. Cell cycle acceleration may be a common strategy used by animals to increase regenerative proliferation. For example, G1 length is reduced during *Drosophila melanogaster* larval midgut regeneration, and G1 and S phases are shortened during axolotl spinal cord regeneration^86,87^. Intriguingly, at least one study has shown that EVs promote cell proliferation by accelerating the G1/S transition^88^. Our observation that EdU-positive neoblasts with S and G2 DNA content increase significantly in EV-injected vs. buffer-injected animals (Fig. 3F) demonstrates that EVs can magnify the proliferative response of a small injury in planarians, and is consistent with acceleration of G2 and/or other cell cycle phases (Fig. 4A). Additionally, although few studies have implicated EVs in cellular division symmetry, EVs could also skew division towards stem cell renewal, rather than differentiation, increasing neoblast numbers. In uninjured planarians, knockdown of the secreted EGF-like ligand Neuregulin 7 increases symmetric neoblast divisions by ∼45%^89^. EVs might play a similar role during regeneration to increase neoblast numbers (Fig. 4B). Future work will be needed to distinguish between cell cycle acceleration and promotion of renewal divisions, and to determine whether these roles are evolutionarily conserved in other animal regeneration models.

**Figure 4.**
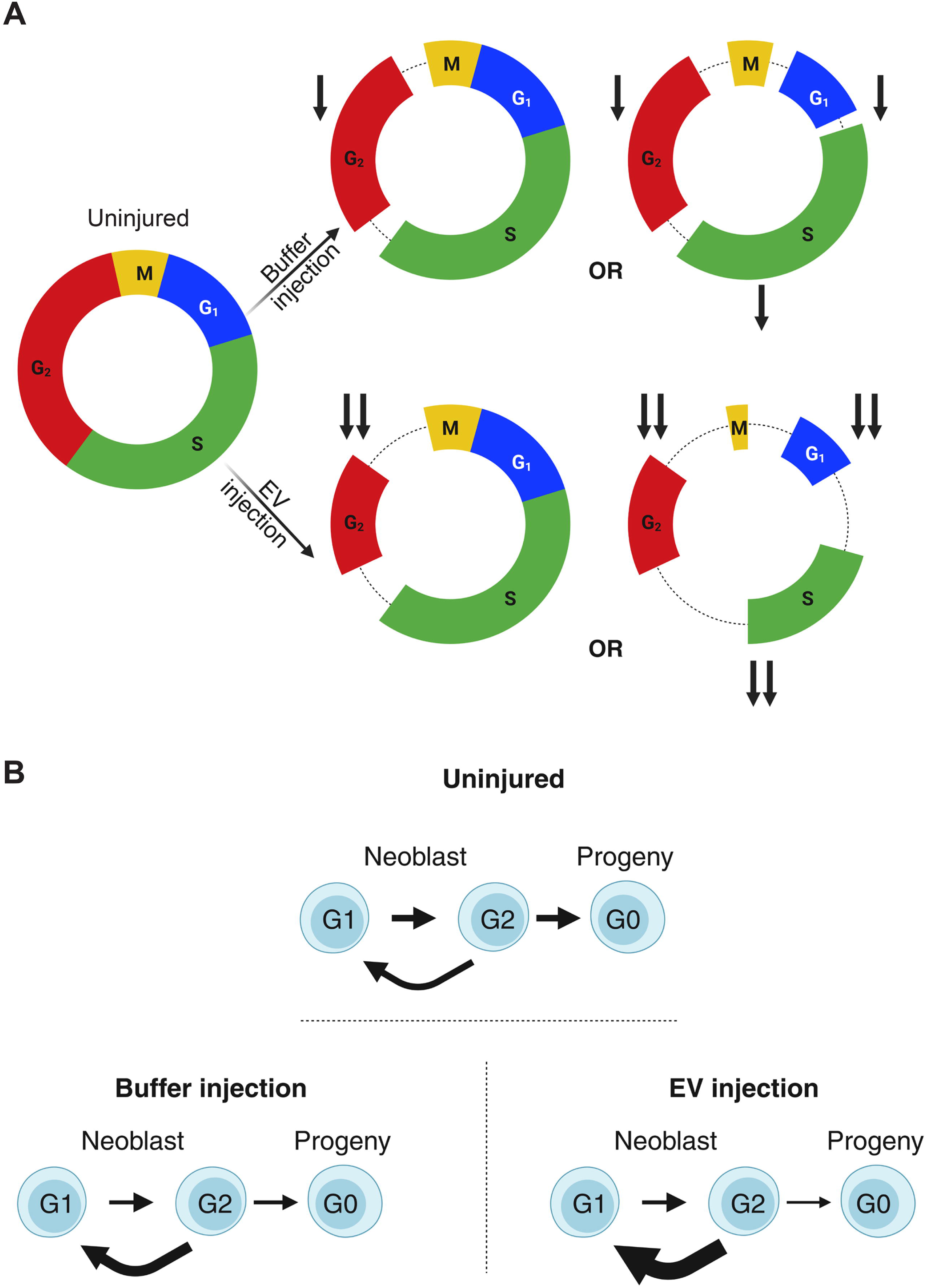
Putative cellular mechanisms of EV-induced proliferation. **(A)** EV injection could decrease the length of neoblast G2 phase, which is thought to occur in response to tissue damage. EV injection could also reduce M, G1, and/or S phase length. Arrows indicate degrees of cell cycle phase shortening. **(B)** EVs may also promote symmetric division, resulting in additional neoblast daughters rather than differentiating progeny. Either or both mechanisms would result in more EdU-positive cells, as we observed. Created in Biorender.com.

EVs could also play additional roles during regeneration, such as modulating cell survival, differentiation, or migration^9,10^. Analysis of EVs enriched at different time points of regeneration, and larger scale characterization of EV protein and nucleic acid cargo, will be needed to identify mechanisms underlying EV function. In addition, separation of EVs into more specific fractions to assess the role of large and small EVs, and development of methods for enriching EVs from distinct cellular sources, will be important next steps to characterize heterogeneity and functions of subclasses of planarian EVs. Importantly, we cannot rule out that co-isolated lipoprotein particles (LPs) contribute to EV-induced proliferation, although previous work suggests that planarian LPs may primarily promote stem cell progeny differentation^22^. Alternative purification strategies will help to clarify this issue. Finally, our observations that cell cycle state and differentiation status influence EV uptake rate suggest that planarians may be useful for understanding EV-target cell interactions during regeneration.

Our study shows that EVs enriched directly from regenerating tissue of an animal capable of whole-body regeneration can promote stem cell proliferation. Thus, the planarian is an amenable animal model to understand how EVs regulate cell behaviors during regeneration, how stem cell responsiveness to complex EV pools is controlled, and how tissue damage influences EV biogenesis. Long term, advances in understanding planarian EV biology should inform and complement efforts to develop EV-based regenerative treatments for human tissue injuries.

## Author Contributions

Priscilla Avalos: Conceptualization, experimental design, new method development, investigation, data collection and analysis, and writing (first draft and editing). Lily Wong: Conceptualization, experimental design, new method development, data collection and analysis, and manuscript editing. David Forsthoefel: Conceptualization, project supervision, funding acquisition, experimental design, bioinformatic analysis, and manuscript editing.

## Acknowledgments

We thank the Forsthoefel Lab for invaluable discussions. We thank Bethany Hannafon and James Lausen (OUHSC Dept. of Pathology) for guidance in EV isolation and characterization. We are grateful to Graham Wiley and the OMRF Clinical Genomics Core; Ben Fowler and Justin Willige in the OMRF Imaging and Histology Core; Jacob Bass in the OMRF Flow Cyotemtry Core; OMRF Gnotobiotic Mouse Core (X-irradiator); Stuart Glenn and OMRF IT/Research Computing Services; and Rajewswari Raguraman, Akhil Srivastava, Ramesh Rajagopal in the OUHSC Stephenson Cancer Center Molecular Imaging Core for expert technical assistance. We thank Gaurav Varshney in the OMRF Genes and Human Disease Research Program for use of the microinjection apparatus.

## Supplementary Figure Legends

**Supplementary Figure 1.**
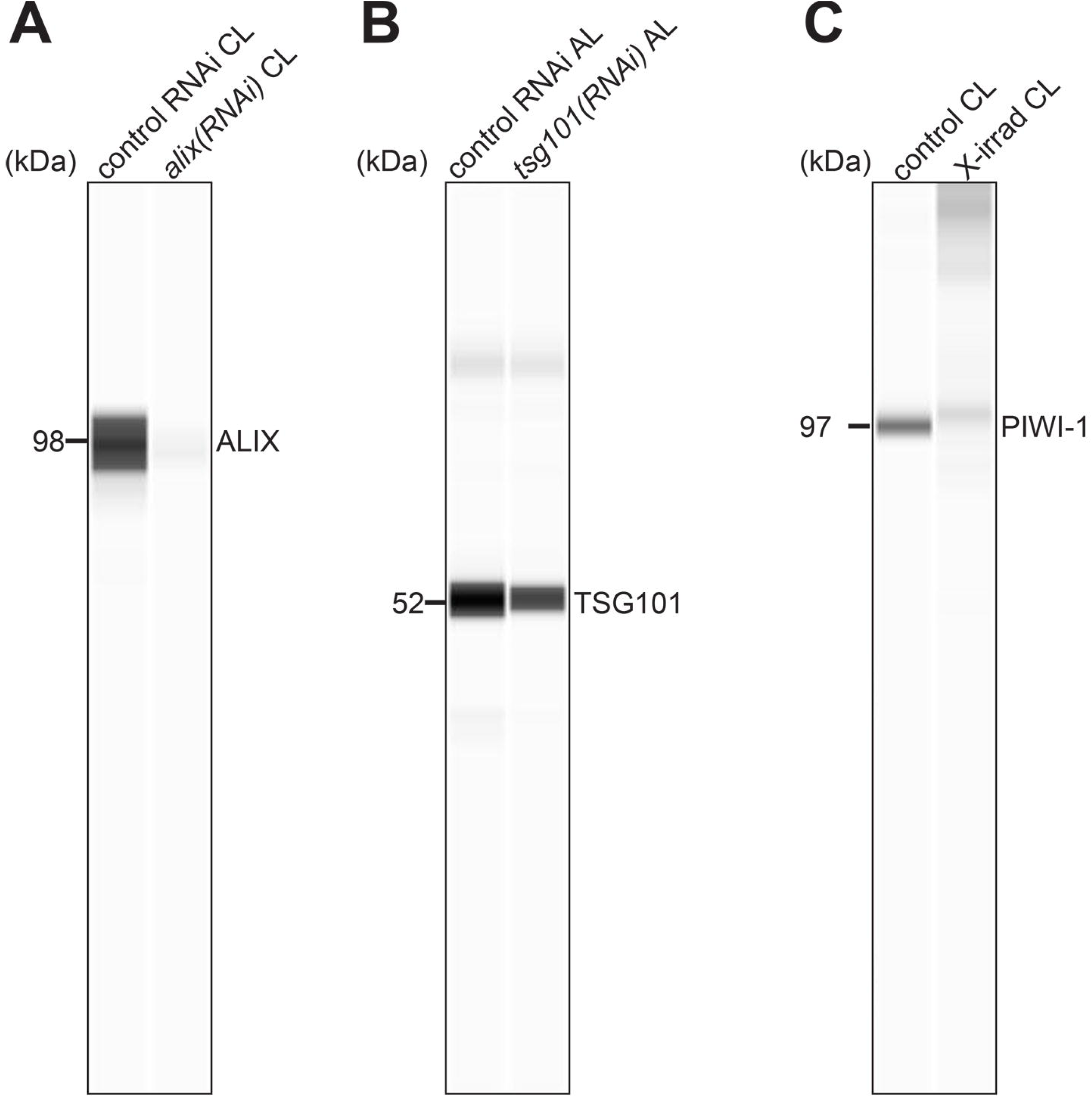
Validation of custom antibodies against ALIX, TSG101, and PIWI-1. **(A-C)** Simple Western of anti-ALIX tested on cell lysates **(A)**, anti-TSG101 tested on animal lysates **(B)**, and anti-PIWI-1 tested on cell lysates **(C)** comparing control (left lanes) vs. knockdown (right lanes) samples (**A-B**) or unirradiated (left lane) vs. irradiated (right lane) samples (**C**), loaded in equal concentrations based on BCA total protein quantification. CL, cell lysate. AL, animal lysate.

**Supplementary Figure 2.**
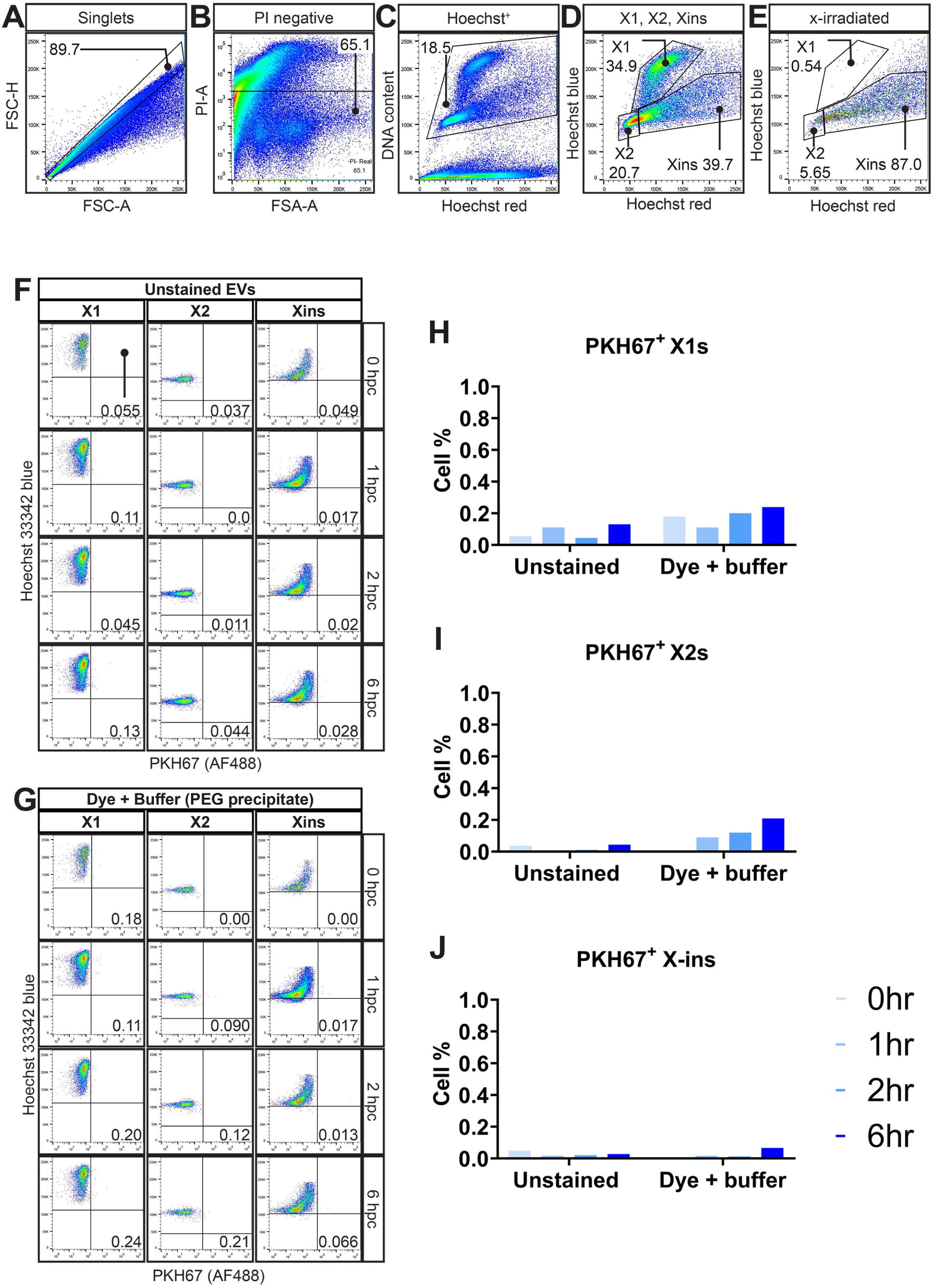
Gating strategy and negative controls for EV uptake experiment. **(A-C)** Representative flow cytometry plots gated to remove doublets **(A)**, dead cells **(B)** and debris **(C)**. **(D-E)** Representative flow cytometry plots showing the gated cell populations distinguished by DNA content into X1, X2, and X-ins of an unirradiated sample **(D)** and an x-irradiated sample 4 days post-irradiation **(E)**. **(F-G)** Representative flow cytometry plots showing the fluorescence intensity of PKH67 (Alexa Fluor 488, x-axis) of X1 (top), X2 (middle), and X-ins (bottom) cells treated with unstained EVs **(F)** or PHK67 plus buffer (PEG precipitate) **(G)** at increasing time points. **(H-J)** Quantification of (**E**) and (**F**) at increasing time points in the X1 (**H**), X2 (**I**), and X-ins (**J**) gates. hpc, hours post-culture.

**Supplementary Figure 3.**
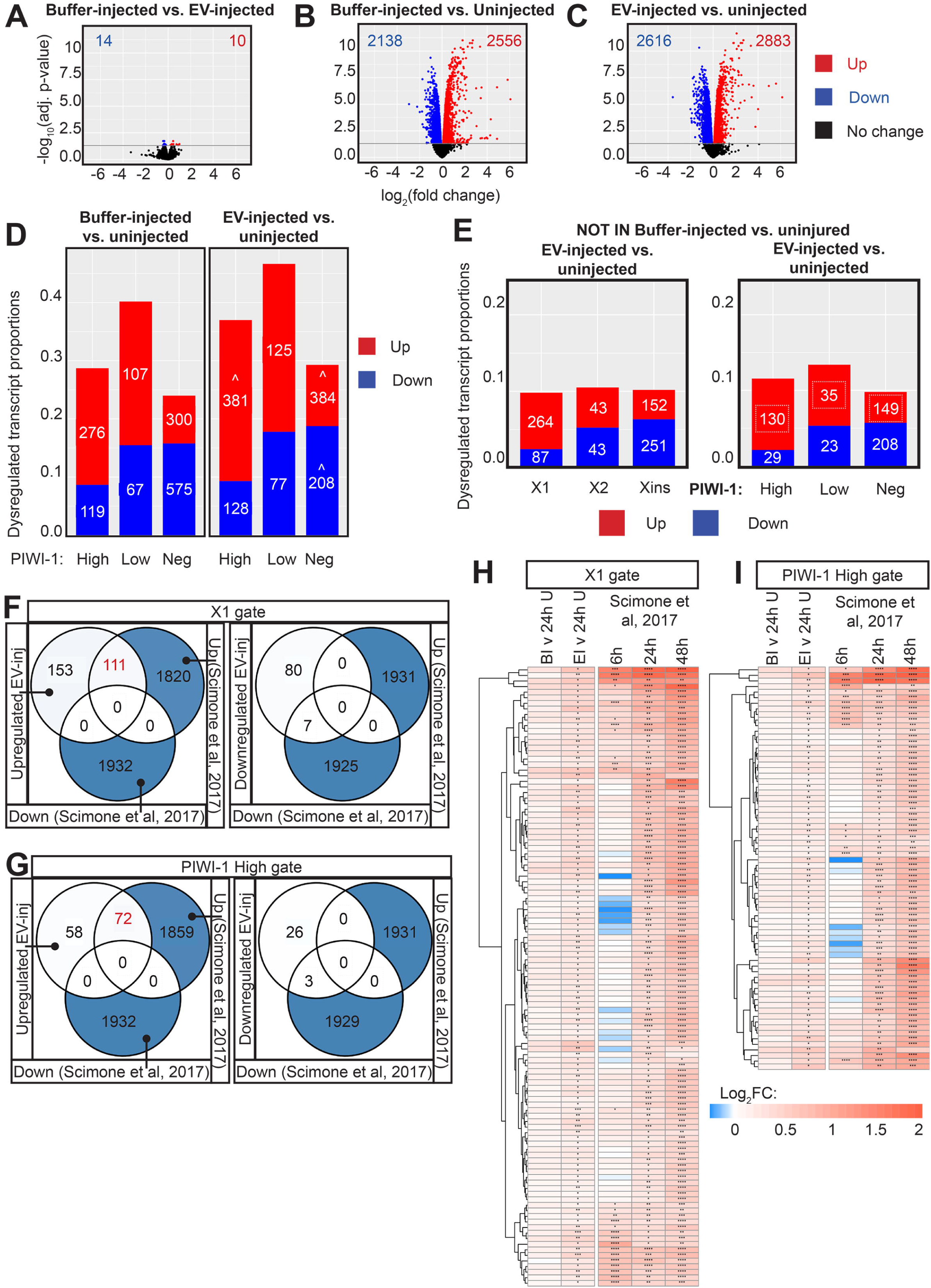
Further analysis of EV-responsive genes in buffer- and EV-injected samples. **(A)** Volcano plots of differentially expressed genes comparing EV-injected vs. buffer-injected **(A),** buffer-injected vs. uninjected **(B),** and EV-injected vs. uninjected **(C)**. Number of significantly up- and down-regulated genes are displayed. **(D)** Number and ratio of PIWI-1-High (1,376), PIWI-1-Low (433), and PIWI-1-Negative (3,646) signature transcripts up- or down-regulated in buffer-injected vs. uninjured (left) and EV-injected vs uninjured (right) animals. Statistically significant differences in transcript number for EV-injected vs. uninjected compared to buffer-injected vs uninjected (i.e., uniquely responsive) are denoted with a caret (^) on the right plot (Fisher’s exact test). **(E)** Number and ratio of X-gate signature transcripts (left) and PIWI-1-gate signature transcripts (right) that are uniquely up- or down-regulated only in EV-injected vs uninjected animals (i.e., not in buffer-injected vs. uninjected). **(F-G)** Venn diagrams showing the overlap of uniquely EV-responsive signature transcripts in X1 (**F**) and PIWI-1-High cells (**G**) with injury-responsive transcripts from 24 hpa blastema tissue (Scimone et al., *Nature,* 2017). Uniquely upregulated X1 and PIWI-High transcripts are shown in red. **(H-I)** Heatmaps of the 111 uniquely upregulated X1 signature transcripts in EV-injected animals **(H)** and 72 uniquely upregulated PIWI-1- High signature transcripts in EV-injected animals **(I)** that are also significantly upregulated in 24 hpa blastemas. Expression changes (log_2_FC) in buffer-injected vs uninjured, EV-injected vs uninjured, and in blastema tissue at 6, 24, and 48 hpa vs. 0 hpa are shown. BI, buffer-injected; EI, EV-injected, U, uninjected; FC, fold change; hpa, hours post-amputation. Asterisks indicate significance (FDR-adjusted p-values) of logFCs: *p<0.05, **p<0.01, ***p<0.001, ****p<0.0001.

**Supplementary Figure 4.**
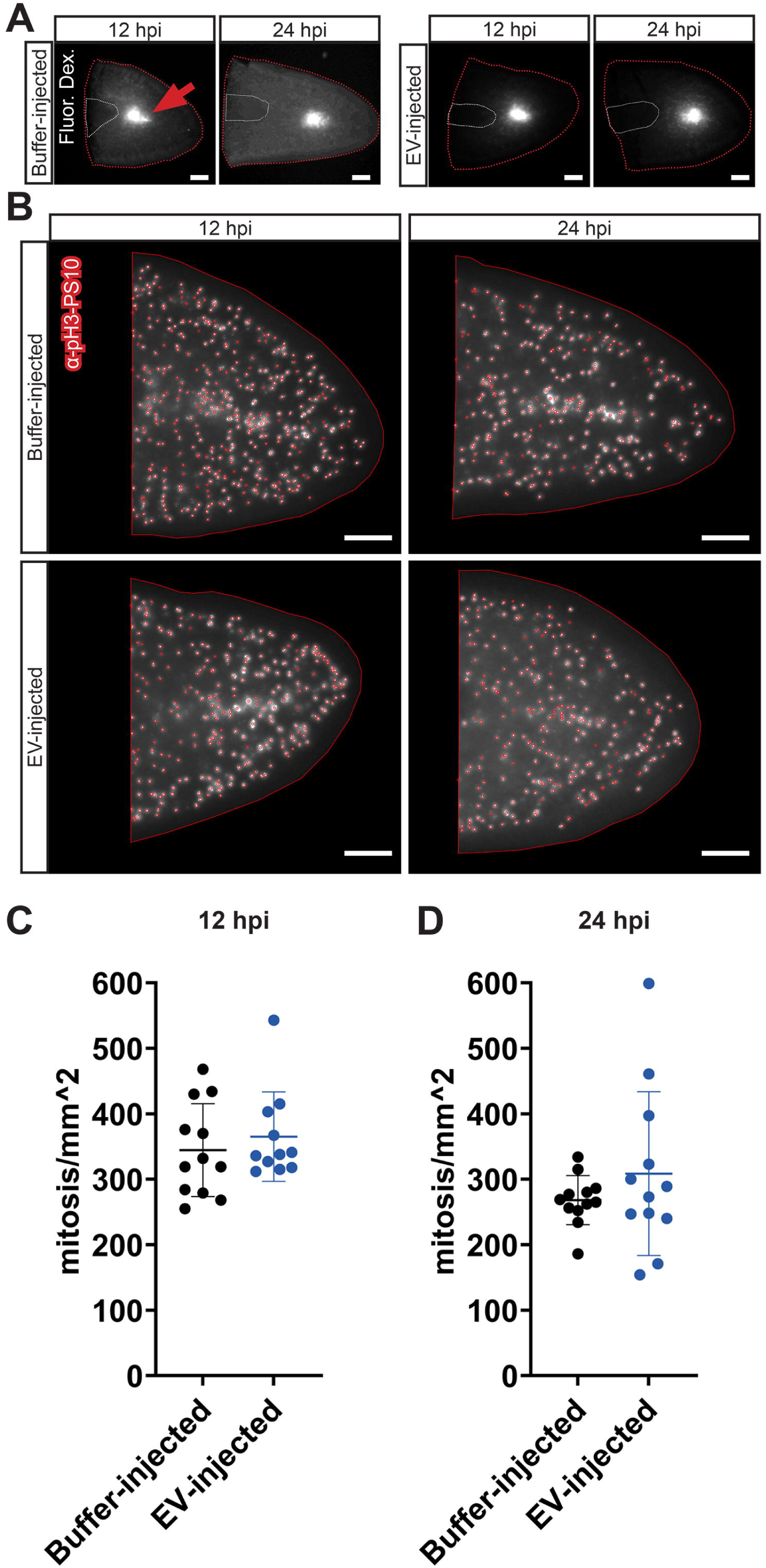
EVs do not significantly increase the number of mitotic cells at 12 and 24 hpi. **(A)** Epiluorescent micrographs of planarian tail fragments (post-fixation) showing the injection site indicated by Alexa-546 fluorescent dextrans. **(B)** Epifluorescent micrographs of mitotic cell labeling with anti-phospho-histone H3- serine10 (PS10) of injected tail fragments. Each region indicated by a red circle (automated detection with FindMaxima in ImageJ) is a PS10^+^ cell nucleus. **(C-D)** Quantification of PS10^+^ cells per mm^2^ in buffer- or EV-injected samples at 12 hpi **(C)** or 24 hpi **(D)**. Unpaired t-tests with Mann Whitney’s correction. Error bars: mean ± S.D., n = 12 (buffer-injected 12 hpi), 11 (EV-injected 12 hpi), 12 (buffer-injected 24 hpi), or 12 (EV-injected 24 hpi) biological replicates per condition. Scale bars: 200 µm (**A** and **B**).

**Supplementary Figure 5.**
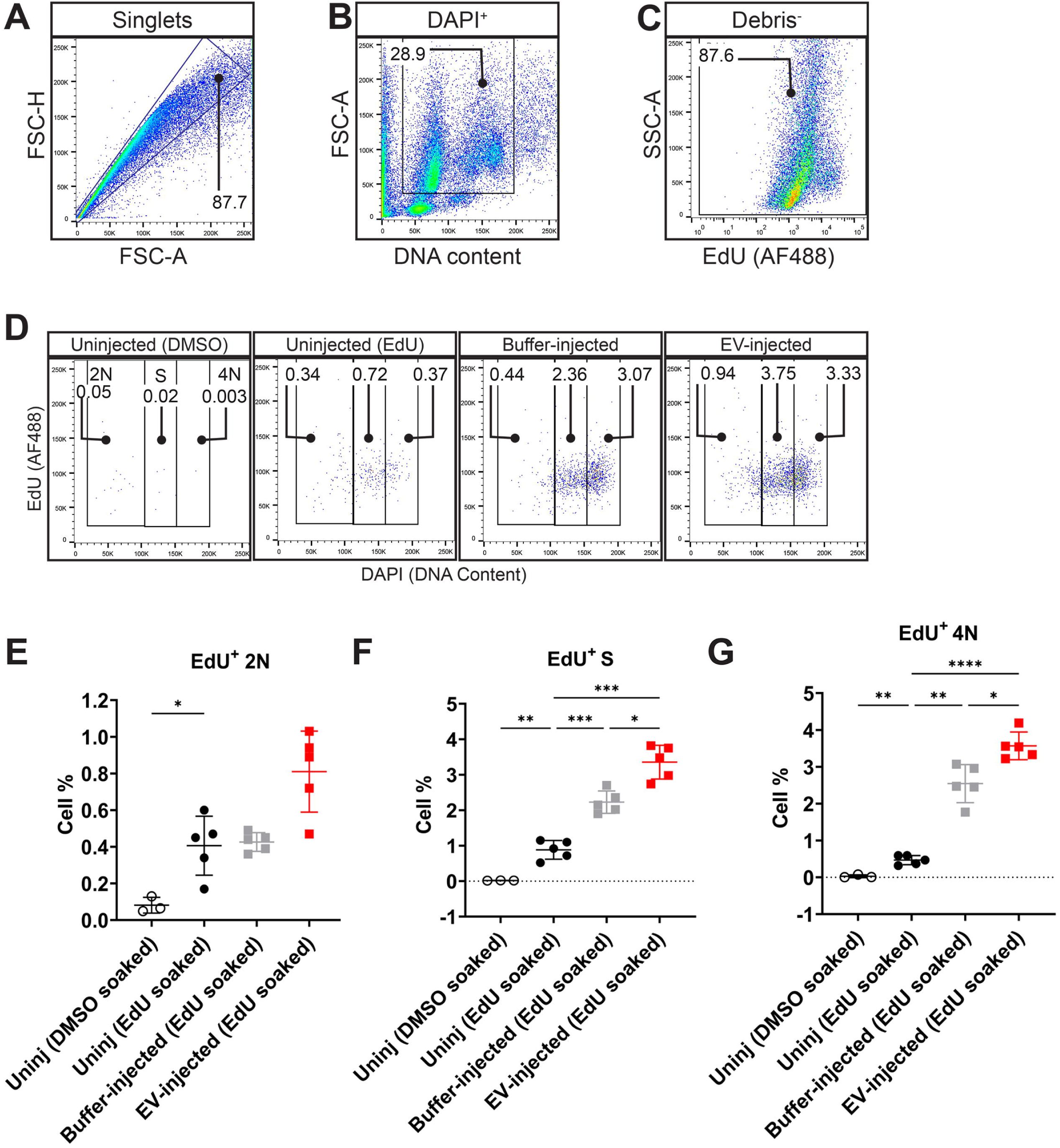
Gating strategy for EdU labeling and DNA content of EdU-positive cells. **(A-C)** Representative flow cytometry plots gated to remove doublets **(A)**, cell aggregates and DAPI negative events **(B),** and debris **(C). (D)** Representative flow cytometry plots showing the fluorescence intensity of EdU^+^ (Alexa Fluor 488, y-axis) cells with 2N, S (>2N but <4N) and 4N DNA content (x-axis). **(E-G)** Quantification of the percent of all DAPI+ cells with 2N **(E)**, S (**F),** and 4N DNA content **(G)** that were also F-*ara-*EdU-positive. Brown-Forsythe and Welch ANOVA test with Dunnett’s T3 multiple comparisons test. Error bars: mean ± S.D., n = 3 (Uninjured DMSO-soaked), 5 (Uninjured EdU-soaked), 5 (buffer-injected), or 5 (EV-injected) biological replicates per condition. (**E**) *p=0.03. (**F**) From left to right: **p=0.008, ***p=0.005, *p=0.0158, ***p=0.0003 (Uninjured vs. EV-injected). **G** From left to right: **p=0.0035, **p=0.0041, *p=0.044, ****p<0.0001.

